# A laboratory method for simulating the effects of subsequent fires on pyrogenic organic matter at different exposure depths in a sand matrix

**DOI:** 10.1101/2024.07.30.605814

**Authors:** Mengmeng Luo, Kara Yedinak, Keith Bourne, Thea Whitman

## Abstract

**Background:** Across a variety of anthropogenic and natural contexts, fire can reoccur in a previously burned location. However, effects of subsequent fire on preexisting pyrogenic organic matter (PyOM) stocks are difficult to discern. Laboratory experiments offer a powerful approach to investigating how subsequent fire impacts the preexisting PyOM.

**Aims:** We aimed to design a highly repeatable laboratory method to effectively measure the impacts of subsequent fires on PyOM at different soil depths while addressing key limitations of previous methods.

**Methods:** Jack pine (*Pinus banksiana* Lamb.) log burns were used to parameterize realistic heat flux profiles. Using a cone calorimeter, these profiles were applied to buried jack pine PyOM to simulate variable reburn fire intensities.

**Key results:** In general, higher heat flux and shallower depths led to more mass loss of PyOM from combustion and more heat exposure.

**Conclusions:** Our reburn method offers a highly replicable way to simulate specific fire scenarios. Conditions that result in more heat exposure (higher heat fluxes, shallower depths) are likely to lead to more loss of PyOM in subsequent fires.

**Implications:** The customizable method could simulate different fire scenarios to investigate spatial variability within a given fire event, or to study the effects of fire on different types of biomass or organisms, such as microbes.

**Summary text:** Our paper illustrates a laboratory method to better quantify loss of preexisting PyOM in soil after a fire.

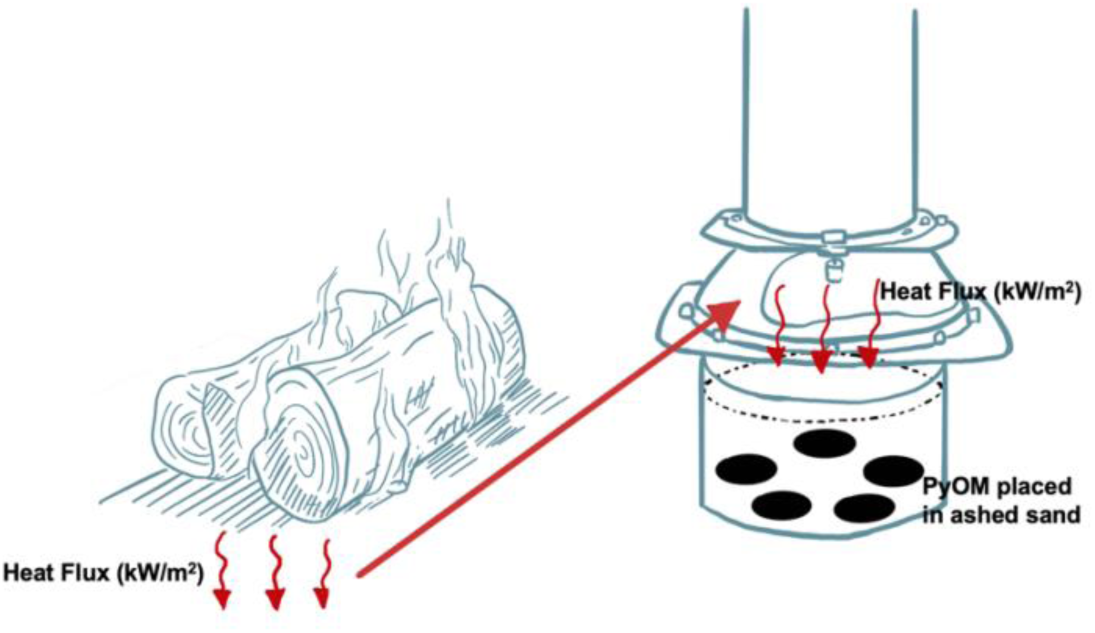

## Introduction

Whether due to naturally high-frequency fire regimes (Holden et al. 2010; Prichard et al. 2017), decreasing fire return intervals (Westerling et. al. 2006; Mansoor et. al. 2022), or repeated prescribed fires used to maintain or restore landscapes (Prichard et al. 2017; Lutz et al. 2020; Saberi and Harvey 2023), subsequent fires (*i.e.*, fires where a previous fire has occurred within ecologically-relevant history) are an important part of understanding the context in which a given fire occurs. Pyrogenic organic matter (PyOM), produced from incomplete combustion of organic materials (Bird et al. 2015), represents a persistent (Spokas 2010) and, in many cases, large component of soil organic carbon (C) (Czimczik et al. 2007; Boot et al. 2014; Reisser et al. 2016). PyOM is generally a larger fraction of soil C in ecosystems with more frequent fires (Reisser et al. 2016) and those for which fuel-reduction efforts include the production and spreading of PyOM for soil health and C sequestration purposes (Franco et al. 2024). However, how subsequent fire affects PyOM produced in previous fires has not been well-studied, despite its importance for our understanding the net effects of fire on C stocks.

Important constraints limit our understanding of these effects. Of the relatively few previous studies that have examined the impact of fires on PyOM, most of them are field studies. For example, the study conducted by Santín et al. in 2013 represents one of the earliest field and lab experiments on the net effect of fire on the pre-existing PyOM (Santín et al. 2013). This study provides direct evidence that fire is an important driver for PyOM loss. Past relevant studies have generally concluded that in natural ecosystems, fire can add, consume, and transform PyOM (Saiz et al. 2014; Tinkham et al. 2016; Doerr et al. 2018; Bartoli et al. 2021). However, constraints in typical methodological designs can be grouped in four key areas: 1) incomplete representation of PyOM, 2) overlooking the PyOM in mineral soil, 3) contamination (*i.e.*, introducing new PyOM) from the fuel, and 4) difficulty in quantifying fire intensity, therefore limiting predictive power or extrapolation beyond the specific system.

### Incomplete representation of PyOM

It is extremely difficult to completely collect or quantify all PyOM in a field setting. Prior studies have found that the particle size and shape of PyOM do not significantly affect mass loss through combustion, within the range of particle sizes studied (Santín et al. 2013; Tinkham et al. 2016; Bartoli et al. 2021). However, these findings reflect the fact that the results from those studies were not based on all the portions of PyOM samples used for the experiments, as smaller size fractions of PyOM were not collected for the analysis. Small-sized PyOM, including soot and aerosol “black carbon” (BC), are often highly persistent but are generally too mobile and/or small to be effectively collected and analyzed (Ansley et al. 2006; Matosziuk et al. 2019), with much of these materials being transported to the atmosphere by wind and deposited outside the system boundary. Consequently, studies including or focusing on the small-size fractions of PyOM are limited, despite the potential implications for C sequestration.

### Overlooking the PyOM buried in mineral soil

A second challenge in studying PyOM after subsequent fires is the difficulty in separating or distinguishing it from other soil organic matter (SOM). One group of methods for isolating PyOM is typically destructive and relies on the general principle of removing chemically labile organic matter (OM) and classifying the rest as PyOM (Zimmerman and Mitra 2017). Thermal oxidation and acid oxidation are two of the most commonly used methods. One example of thermal oxidation is CTO375, which removes SOM and leaves BC by heating the samples at 375°C in the presence of oxygen for over 20 hours (Gustafsson et al. 1996; Hatten et al. 2008). Benzenepolycarboxylic acid (BPCA) is a representative example of acid oxidation. It uses strong acids (like HCl) to dissolve the SOM, leaving aromatic compounds (BPCA) as chemical markers for condensed aromatic C (Brodowski et al. 2005; Matosziuk et al. 2020). Both kinds of methods infer PyOM from compounds with high chemical recalcitrance, which is not fully accurate because the highly degradable portion of PyOM is ignored (‘false negatives’), and some of the measured aromatic C may not have been produced from heating (‘false positives’).

Typical non-destructive methods, such as mid-infrared (MIR) spectroscopy also suffer from potential false positives and false negatives, and identify PyOM without separating it from other SOM (Zimmerman and Mitra 2017), thereby precluding direct chemical and biological analyses of the PyOM on its own. As a result of these methodological limitations, most studies have focused on the PyOM on the surface or in the organic horizon, where it can be visually separated from unburned OM, and not in the mineral soil. However, over time, surface PyOM moves from the organic horizon to mineral soil (Tinkham et al. 2016; Matosziuk et al. 2020). This vertical movement of PyOM can occur through bioturbation or translocation with water, while PyOM can also be produced at depth through the charring of roots and buried biomass *in situ* (Hobley 2019; Soucémarianadin et al. 2019). Previous studies of subsequent fires have suggested that PyOM residing in deeper soil horizons can be protected from heat and combustion due to soil insulation (Santín et al. 2013; Saiz et al. 2014; Doerr et al. 2018; Bartoli et al. 2021), meaning it is particularly important to understand the interactions between burial depth and PyOM loss or alteration in subsequent fires. However, there is a lack of comparative experiments to validate these expectations, with most studies focusing on PyOM either on the surface or within the litter layer, not in the mineral soil.

### Contamination from fuel

The use of a fuel bed, typically composed of woody materials, is a common method for sustaining heat during the PyOM burning process, designed to provide a realistic scenario for a forest fire. However, this method can contaminate the preexisting PyOM in the system due to the addition of new PyOM from the fuel, making it challenging to analyze the net effect of fire on the PyOM. Some studies have attempted to address this issue by wrapping the PyOM sample in a metal mesh bag to separate it from the fuel (Santín et al. 2013; Bartoli et al. 2021), but the metal can alter the heat transfer to the sample, and small particles can still leak out from or into the mesh bag.

### Quantifying fire intensity

Fire intensity is defined as the ‘energy output’ of a fire and is not explicitly linked to any effects caused by fire (Keeley 2009). However, quantifying fire intensity, particularly in natural settings, is difficult. First, quantitative measurements of fire intensity can be misused or vaguely defined. Temperature is a useful indicator for how much heat is dosed to the samples, (Santín et al. 2013), but is not as direct and accurate as energy fluxes. Fireline intensity (with the unit kW m^-1^) has also been used as a proxy for fire intensity in some studies (Doerr et al. 2018). However, this concept cannot be easily translated to laboratory settings, as fireline intensity is commonly used for scenarios where there are moving flames (Keeley 2009). So, it does not accurately quantify the energy released from glowing or smoldering fuels, which may last for hours after the flame goes out (Keeley 2009; Kremens et al. 2010). Second, in laboratory settings, the simulated fires in the past studies are not always tied to real-world parameters (including temperature, duration, and direction).

### Methodological approach and hypotheses

We sought to develop a method for simulating the effects of subsequent fire on PyOM that effectively addresses each of the limitations discussed above. We developed lab simulations of subsequent fire using a cone calorimeter with PyOM in a sand matrix, with the following characteristics: 1) All portions of PyOM after the reburn are collected and analyzed. 2) PyOM is buried at a range of depths within a sand matrix, to include the overlooked mineral soil while also ensuring that the PyOM sample is the only C input in the system. 3) Instead of applying a fuel bed to sustain the heat for the burn, we apply a precise heat flux (HF) dose using a cone calorimeter. 4) The HF profiles, representing high and low fire intensity, were drawn from the data captured from HF sensors under tree log burns to parameterize a realistic fire scenario.

We hypothesized that higher HF and shallower depths would have more PyOM mass losses. At the same HF, we predicted that PyOM on the surface would be subject to more mass loss than PyOM at both 1 cm and 5 cm depths. At the same depth, we predicted that high HF fire would consume more PyOM than low HF fire.

## Methods

### Ecosystem, site description, and experimental overview

We designed our burn experiments to generally reflect the ecosystem of the jack pine (*Pinus banksiana* Lamb.) barrens in Wisconsin (WI), where the soil is characterized as coarse sands with low nutrients, low water holding capacity, and prone to drought (Radeloff et. al. 1999, 2004). Jack pines are known for their dependence on fire to open their cones for seed release and germination (serotiny). In the Northwestern Pine Barren of WI, the stand-level serotiny gradually decreases from the north to the south (Radeloff et al. 2004). In the southmost region, jack pine standings are subject to the lowest stand-level serotiny (<24%) due to non-lethal high-frequency fire (with a 3∼7-year fire return interval) and thick tree bark. For this experiment, we used two jack pine trees that were cut down from Wilson State Forest Nursery (43.1461124, -90.6950388) and Hancock Agricultural Research Station (44.122667, -89.530740) in Wisconsin. (Because the locations of these trees are south of most native WI pine barrens and in managed land, we do not expect serotiny to necessarily occur in the trees.)

Briefly, to simulate the effects of subsequent fire on PyOM, we pyrolyzed jack pine wood to produce PyOM, and then burned the resulting PyOM under a cone calorimeter at the Forest Products Laboratory of USDA Forest Service in a sand matrix. We designed the fire to be representative of logs burning on the soil surface in a jack pine stand, including a higher and lower fire intensity. We directly quantified the HF transferred downwards into the soil in this system, and chose this approach to represent a scenario where the greatest heat transfer to soil might be likely to occur, thereby capturing the upper bounds of fire effects in this system.

### Production of PyOM

The PyOM was produced from ground jack pine wood (<2 mm) at a highest treatment temperature of 350°C in a muffle furnace (Thermo Fisher Scientific 1100°C Box Furnace BF51800 Series; Güereña et al. 2015; Zeba et al. 2022) modified to operate under an argon gas atmosphere. 350°C is within the temperature range of a typical low-intensity forest fire (Santín et al. 2016b). We increased the temperature from 25°C to 250°C at 5°C min^-1^, then increased it to 350°C at 5°C min^-1^ with the rotor on (to stir the PyOM in the chamber) and sustained it at 350°C for another 30 minutes with the rotor on. After that, we stopped the rotor, and water was circulated outside the pyrolysis chamber to rapidly cool the PyOM. The PyOM was collected once it reached room temperature. The PyOM yield was ∼35% of the unburned jack pine wood by dry mass. This PyOM thus represents PyOM produced in the first fire, while the fire simulated in the experimental burns with the calorimeter is considered to be the subsequent fire for the purposes of this experiment.

### Determining fire intensity via heat transfer sensing in log burns

In order to ensure the heat treatments were ecologically realistic, we designed experimental log burns to determine the HF dose for the High- and Low-HF treatments. The logs used in this experiment were stored in a drying room at 32°C and 30% relative humidity for six months. The logs were then oven dried at 105°C for >48 hours before the burn. A sand bed was prepared and leveled for the burn with three embedded water-cooled HF sensors. Two were Hukseflux combination Gardon and Schmidt-Boelter heat flux sensors with a measurement range of 0-200 kW m^-^² (model SBG01-200). The third was a Medtherm 64 series Schmidt-Boelter heat flux sensor with a range of 0-114 kW m^-^² (model 64-10SB-18). All the sensors were calibrated under the cone calorimeter before the deployment. The tops of the sensors were at the same level as the surface of the sand bed, and they were linearly aligned in the center of the sand bed (Fig. 1a). One treated log (i.e., small groove cut by hand to increase surface area for combustion), and one untreated log were used for each burn (Fig. 1a). The treated log was oriented with groove facing down and subjected to four natural gas Bunsen burners, located 12 cm below the suspended log, for 10 minutes (Fig. 1b). The treated log was then placed parallel to the untreated log with groove facing the heat flux sensors and angled such that one end was elevated approximately 10 cm above the sand to promote buoyant gas flows. The two logs (treated and untreated) were situated on the sand bed such that the three heat flux sensors were in the middle of the two logs and ran parallel to the length of the logs (Fig. 1a, Fig. 1c, and Fig. 1d.). The principle for this experiment was to sustain a controlled, laboratory-scale burn in order to obtain downward heat flux profiles from the paired logs that would then be input into the calorimeter for burning the PyOM samples.

**Figure 1.**
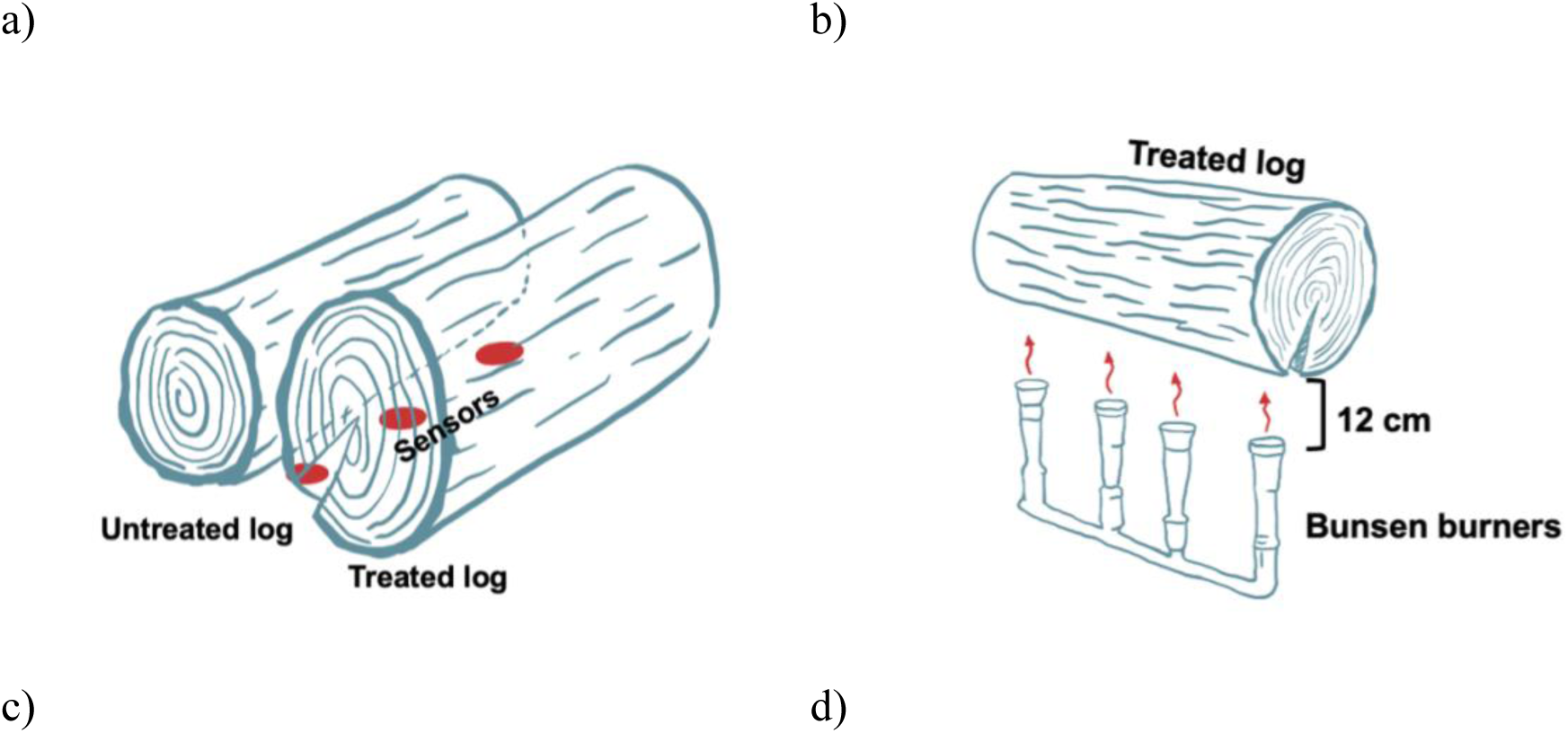
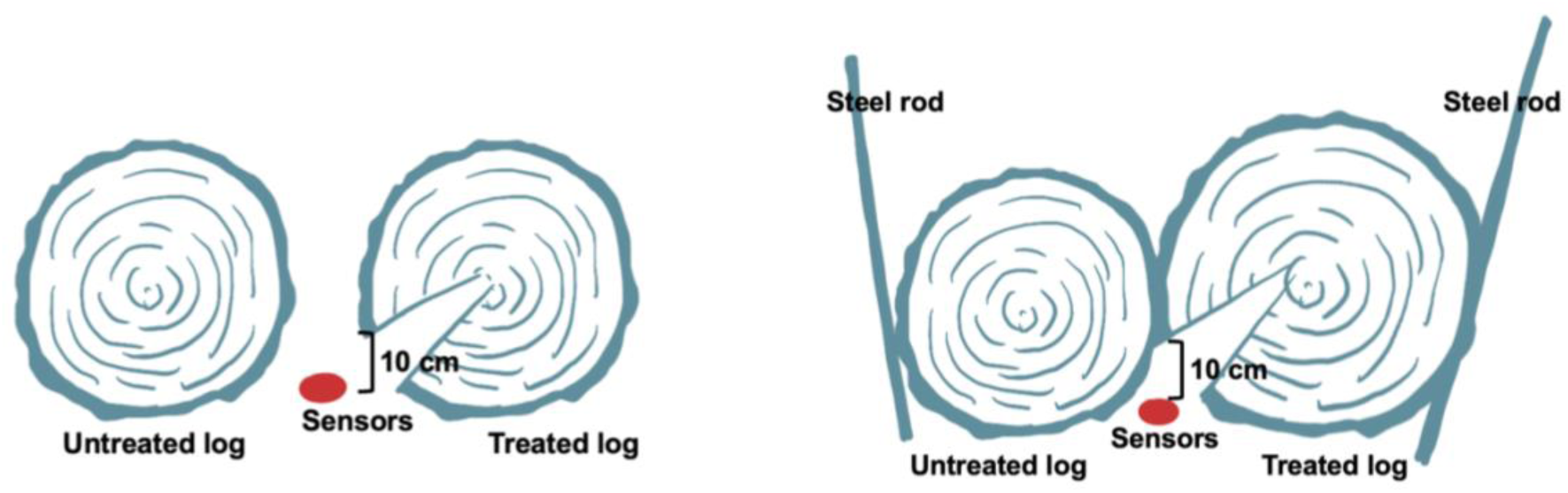
Log burns preparation and setup (corresponding photos displayed in Supplemental Figure S1). a) Logs and sensors’ placement; b) Treated log on burners; c) the treated (right) and untreated (left) logs for capturing Low HF profile (cross section displayed); d) the treated (right) and untreated (left) logs for capturing High HF profile (cross section displayed)

We used the HF data of two 2-log experimental burns to inform two fire scenarios – High HF and Low HF – which represented the upper and lower heat flux profiles out of all the experimental trials we did. Both HF profiles were determined from the three sensors between the two logs. For the Low-HF profile, the treated log was of a similar size to the untreated log, and the two logs were not stabilized in place (Fig. 1c). As the biomass was consumed by fire, the distance between the logs got wider and measured HF declined. For the High-HF profile, we wanted to model higher fuel connectivity and a higher fuel load. Thus, we maintained a relatively constant distance between the two logs with four steel rods that kept the two logs in place (Fig. 1d), and used a treated log that was bigger than the untreated log.

The High-HF profile, representing the ‘high-intensity fire’, was then replicated and monitored in the cone calorimeter using LabVIEW (National Instruments 2013) with a peak HF at 85 kW m^-2^ and a duration of 17000 seconds; the Low-HF profile, representing the ‘low-intensity fire’, was modeled with a peak HF at 45 kW m^-2^ and a duration of 9000 seconds (Fig. 2).

**Figure 2.**
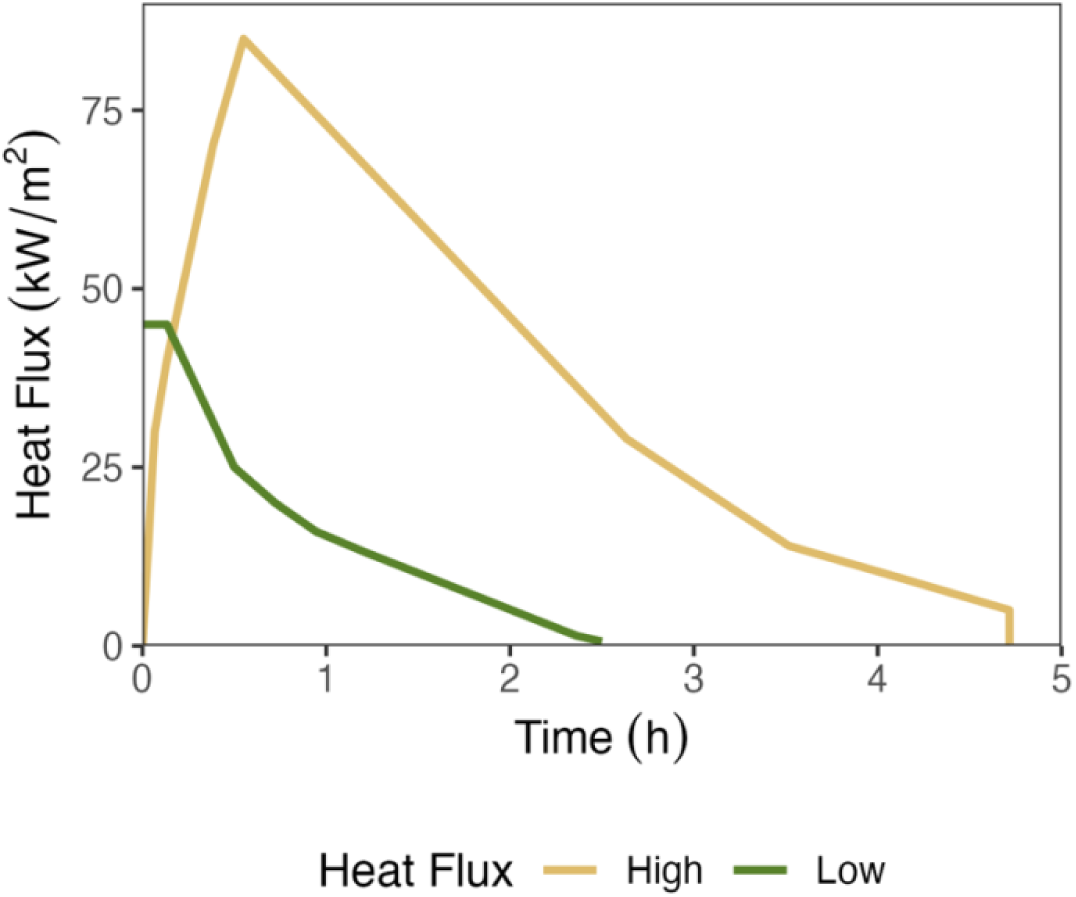
High- and low-HF profiles dosed to the samples and based on log burns

### Sample treatments and the burning matrix

Three different depths of PyOM (Surface, 1 cm, and 5 cm) were tested, and three fire treatments (High HF, Low HF, and Control), were applied. These were combined in a full-factorial design, for nine treatments in total, with five replicates each of 1 g PyOM. AquaQuartz Pool Filter Sand (0.45-0.55 mm) was used as the matrix for all the treatments, representing sandy soils typical of jack pine stands while ensuring a C- and nutrient-free system. The sand was treated in advance to further eliminate any traces of C by heating it at 500°C for three hours in a muffle furnace (‘ashed’). Then it was washed with Milli-Q water to flush out any soluble chemicals, oven dried at 100°C for more than 48 hours, and stored in autoclaved media bottles.

A burn unit was designed – a steel container filled with the ashed and washed sand. The container was modified from a stainless-steel beaker (McMASTER-CARR Stainless Steel Beaker with Handle, 2850 mL capacity, 152.4 mm in diameter and 90 mm in depth). Five PyOM replicates for each fire treatment were placed in the matrix using a custom-designed 3D-printed PLA (polylactic acid filament) sample placer (modeled with Tinkercad; printed with Ultimaker S5, University of Wisconsin-Madison Makerspace), which is a flat plate embedded with five identical open-ended tubes (Fig. 3). Each tube has a diameter of 30 mm and a depth of 16 mm. The center of each tube is equidistant (45 mm) to the center of the plate and the same distance (52.9 mm) from the adjacent tubes (Fig. 3). The sample placer ensured that the samples were placed at the same level in the burn unit and same distance from each other at each treatment.

**Figure 3.**
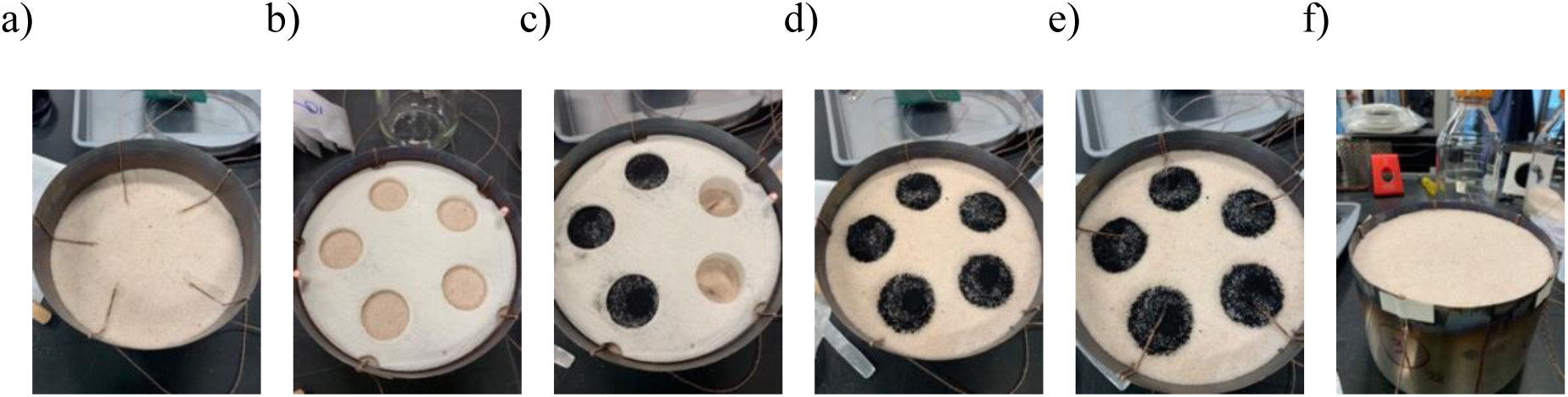
Sample placer (image generated from Tinkercad)

To place the samples at different depths for each treatment, we leveled the sand and pressed the sample placer into the sand. Then we scooped out the sand from each tube and filled the hollows with the samples, and gently removed the sample placer. In order to better simulate field conditions, we mixed the 1 g of PyOM with 8 g of sand before burying it. For the 1-cm and 5-cm depths, we added sand to the top of the samples and leveled the top of the matrix at 10 mm from the top of the container. Thermocouples made from 30-gauge type K wire (Omega Engineering, GG-K-30-SLE) were placed above and below each sample (Fig. 4a-f).

**Figure 4.**
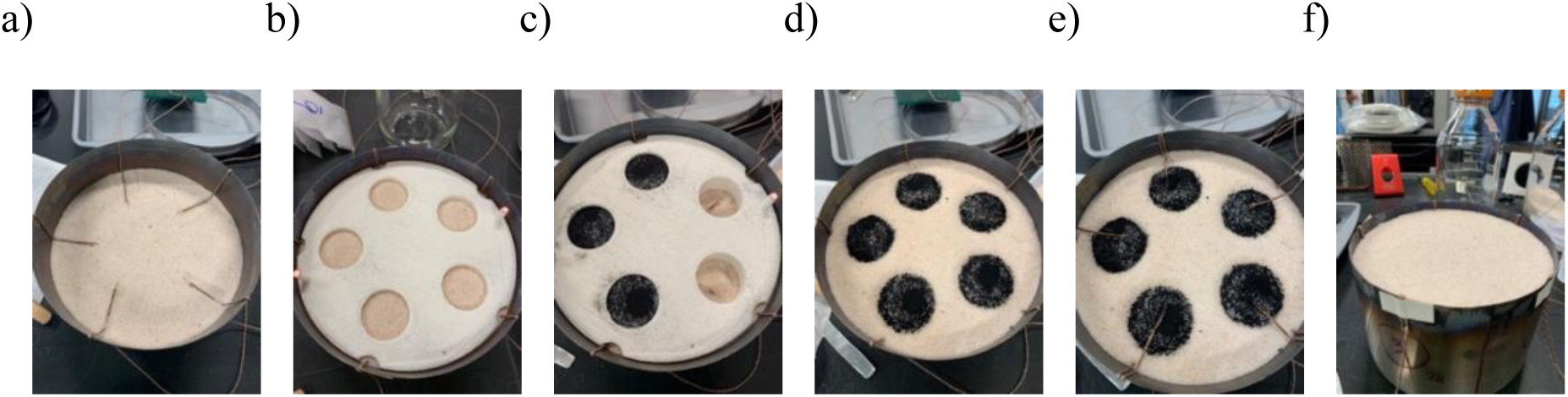
An example of processes for placing the samples for 1 cm-depth treatments. a) Thermocouple wires are placed ∼16 mm below the designated depth, which is the depth of the sample placer tube; b) Sand is filled to the designated depth and press the sample placer in the sand; c) Sand is scooped out from the tubes, which are refilled with the PyOM-sand samples; d) The sample placer is lifted out with the samples remaining in the sand; e) A second set of thermocouple wires are placed on top of the samples; f) The container is filled with sand to bury the samples (this step is skipped for surface treatments).

Everything in the burn unit, including the steel container, ring for stabilizing thermocouples, and thermocouples, were weighed individually and all together before and after the burn to determine total mass loss of the samples and any extra mass loss of the appliances for each treatment. We also ran a “blank trial” for both HF profiles with only sand to get how much sand was volatized during the burn. The mass loss was measured for all the samples in the same burn unit for the same treatment, not individual sample. Mass loss fraction and mass remaining (%) were calculated with Equations 1 and 2:

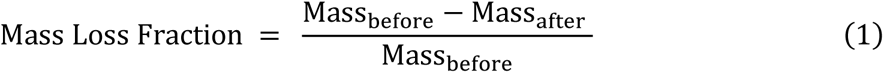

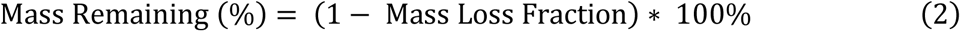

Before placing the matrix under the cone calorimeter, an insulation sheet was wired around the container (Fig. 5) to prevent the thermocouples from being burned and to help ensure energy only enters or leaves the burn unit through the top surface. The burn unit was dosed using the pre-set HF profiles as described above. The temperature data for each thermocouple was recorded using LabVIEW (National Instruments, 2013).

**Figure 5.**
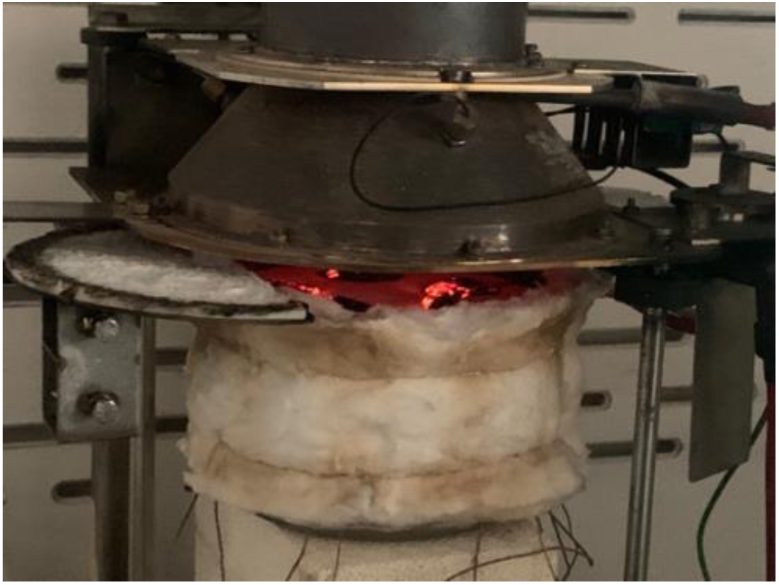
Sample matrix with insulation sheet under the cone calorimeter

After each burn, the burn unit with the sand matrix was carefully removed from the calorimeter and set in a hood to cool. After the temperature of the samples dropped below 200°C, we slid a water-circulating ring onto the burn unit to speed up the cooling process and ensure the cooling was even for each replicate (Fig. 6). After the temperature dropped below 80°C, we took the unit out for sample collection. Each individual sample was collected and reserved for further chemical and biological analyses not described in this paper.

**Figure 6.**
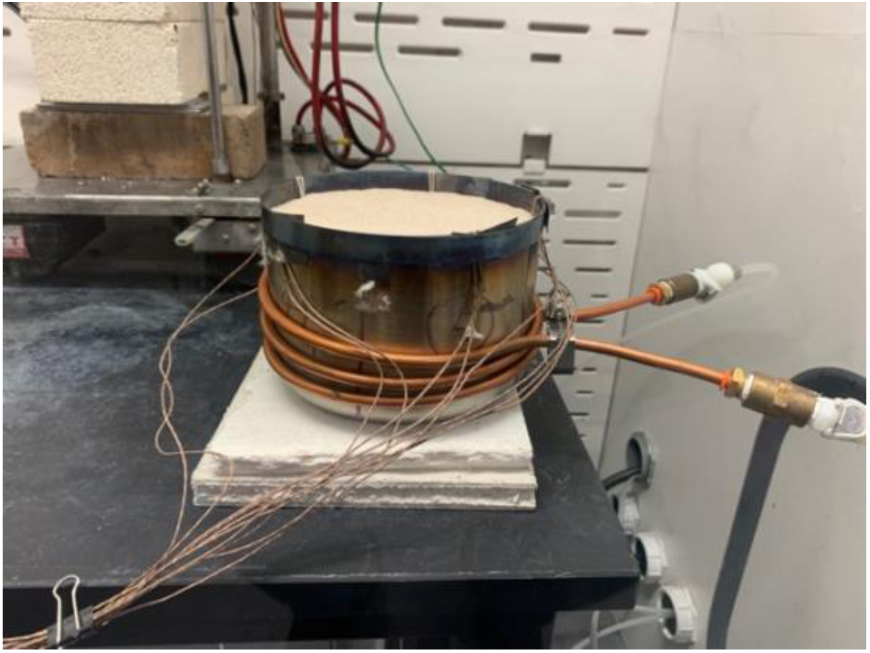
Cooling the samples. Burn unit with sand-PyOM matrix is wrapped in water-circulating ring. Thermocouple wires can be seen attached to the burn unit.

The cone calorimeter in our experiments was under temperature (T) control mode during the burning of the samples (it has both T control and HF control modes). T control mode directly controls the temperature of the cone above the burn unit, while under HF control, it controls the emission of heat based on the HF that the sensor receives, which requires that the sensor be set at the same location as the samples. However, there was not enough space to place the HF sensor and the samples in the same burn unit. Additionally, since the controlling HF sensor is water-circulated for cooling, it could also cool down the samples if they were placed together. Thus, to ensure that we were able to administer the desired HF during the sample burns, we ran a blank burn for each HF treatment, where only the HF sensors were embedded in the burn unit (no samples), in HF control mode while we also recorded the temperature. Thus, we were able to run the calorimeter in temperature control mode during the experimental burns to achieve the same targeted HF. *Statistical analysis*

Calculations and statistical analyses were done using Excel and R (R Core Team, 2022). Figures were created using ggplot 2 (Wickham 2016) in R (R Core Team, 2022). A two-way ANOVA with an interaction term for heat flux and burial depth and Tukey’s HSD (Tukey, 1949; Graves et al. 2019) were used to determine if there are significant differences between treatments. Degree hours, calculated from the area under the temperature profile, were used to quantify the temperature and duration of heat exposure. A binomial logistic model (with function “glm”) (R Core Team, 2022) was used to fit the degree hours and peak temperature as predictors for the fractional mass loss of PyOM.

## Results

### Qualitative observations

PyOM samples in surface treatments under both HF profiles started to combust in the first 5 minutes. During the High HF + 1 cm treatment, smoke escaped from the burn unit within the first 17 minutes from the start of exposure. Thus, we expect that for those three treatments (High HF + Surface, High HF + 1cm, and Low HF + Surface), the biggest chemical changes and PyOM losses might occur much sooner than the full HF profile’s duration. (We were not able to directly track mass loss dynamically during the heating process due to oscillation caused by buoyancy-driven ambient airflow.) Those three treatments also resulted in little visible PyOM left. No obvious smoke was observed for High HF + 5 cm, low HF +1 cm, or low HF + 5 cm.

### Temperature profiles are consistent within and distinctive across treatments

Temperature profiles were consistent across replicates and show distinctive patterns for different HF treatments (Fig. 7). Overall, temperatures decreased with burial depth, and were higher in the High-HF treatments at the same depth (Table 1). The peak temperatures reached during the experiment had a time lag from the peak heat flux emitted from the cone calorimeter. The temperature profile and peak temperature of the High HF + 5 cm were similar to those of Low HF + 1 cm treatment (Fig. 7). Peak temperatures were not significantly different between these two treatments but were significantly different among other treatments (p < 0.001, ANOVA in Table S1, Tukey’s HSD, Table 1). The degree hours were significantly different across each treatment (p < 0.001, ANOVA in Table S2, Tukey’s HSD, Table 1).

**Figure 7.**
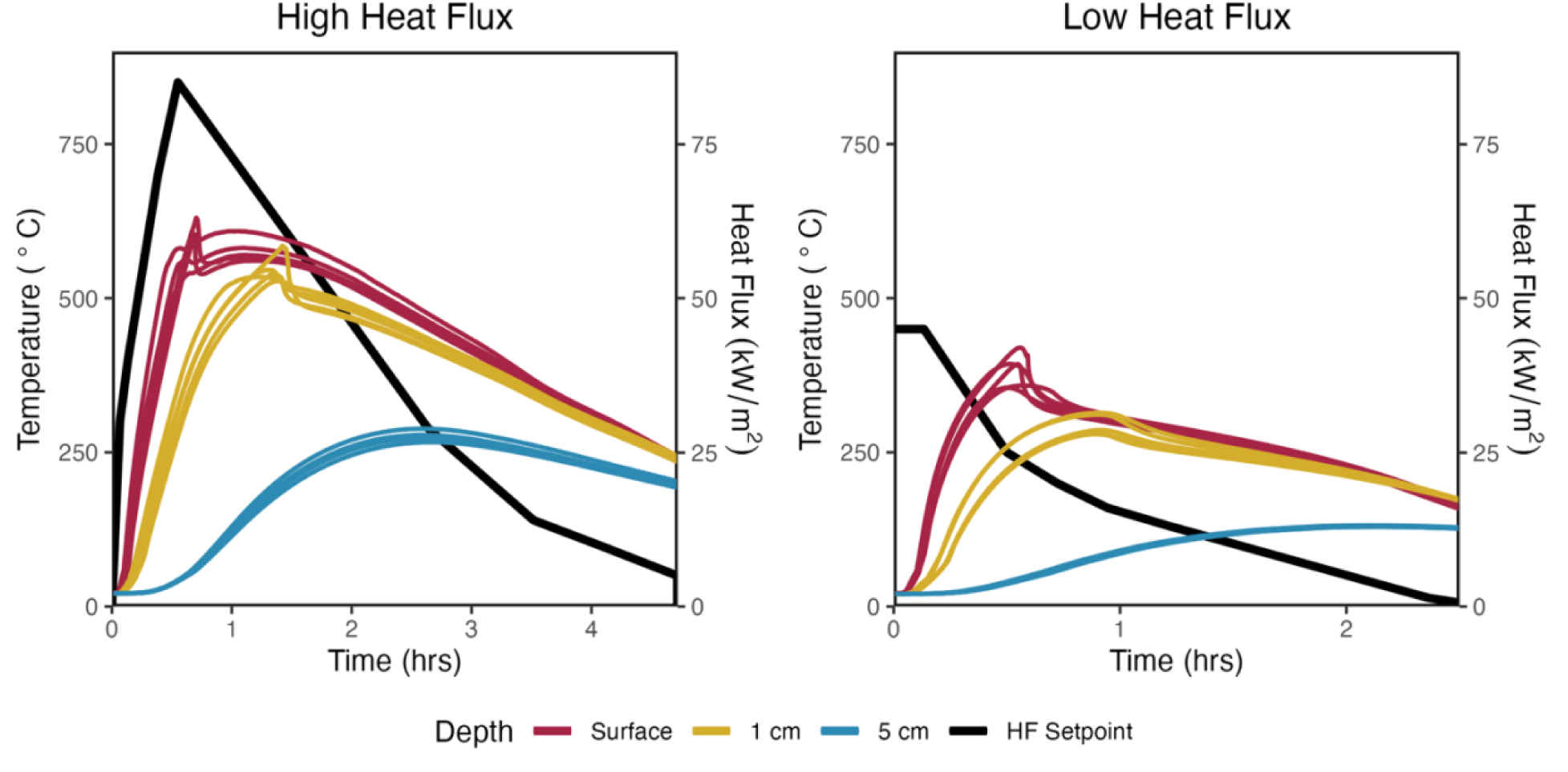
Temperature profiles for the bottom thermocouples for five replicate samples in each HF treatment (colored lines) and the corresponding heat flux profiles (black lines), for the High HF (left) and Low HF (right). The temperature data captured by the top thermocouples were not graphed due to excessive oscillations in the near-surface treatment as the thermocouples became exposed to the air (for full temperature profiles, refer to Supplemental Figure S2). Note different time scales on x-axis. Samples continued to cool to room temperature beyond data plotted here.

**Table 1.** Degree hours and peak temperature data and statistical results (n=5). Different superscript letters (ANOVA, Tukey’s HSD) indicate significant differences across all treatments.

| *Heat Flux Profiles* | **High** |  |  | **Low** |  |  |
| --- | --- | --- | --- | --- | --- | --- |
| *Heating Duration (hrs)* | **4.72** |  |  | **2.5** |  |  |
| *Exposure Depth* | **Surface** | **1 cm** | **5cm** | **Surface** | **1 cm** | **5 cm** |
| Degree hours ($^{\circ}\text{C} \times \text{hrs}$) | 2006.46<br>(57.67)<sup>a</sup> | 1750.15<br>(49.32)<sup>b</sup> | 928.27<br>(26.98)<sup>c</sup> | 639.10<br>(10.90)<sup>d</sup> | 539.47<br>(24.81)<sup>e</sup> | 224.20<br>(3.04)<sup>f</sup> |
| Peak Temperature ($^{\circ}\text{C}$) | 598.91<br>(23.64)<sup>a</sup> | 546.39<br>(22.24)<sup>b</sup> | 276.08<br>(8.03)<sup>d</sup> | 383.85<br>(27.16)<sup>c</sup> | 294.64<br>(16.66)<sup>d</sup> | 130.24<br>(0.94)<sup>e</sup> |

### Greater heat exposure leads to higher PyOM mass loss

Mass loss ranged from almost 100% (High HF + Surface and High HF + 1 cm, 98.86% and 98.45%, respectively) to as little as 6.6% (Low HF + 5 cm) (Fig. 8). Mass loss for High HF + 5 cm, Low HF + Surface, and Low HF + 1 cm were 29.19%, 93.51%, and 47.72%, respectively (Fig. 8). (Note that the entire burn unit was weighed before and after the burn, so we report only the total mass loss of each treatment instead of individual samples.) Peak temperature during the burn was more strongly associated with mass loss than degree hours based on model fitness (indicated by pseudo-R^2^ and AIC; Fig. 9). Based on the right panel of Fig. 9, greater heat exposure generally led to higher PyOM mass loss in an estimated temperature range of 200°C to 400°C

**Figure 8.**
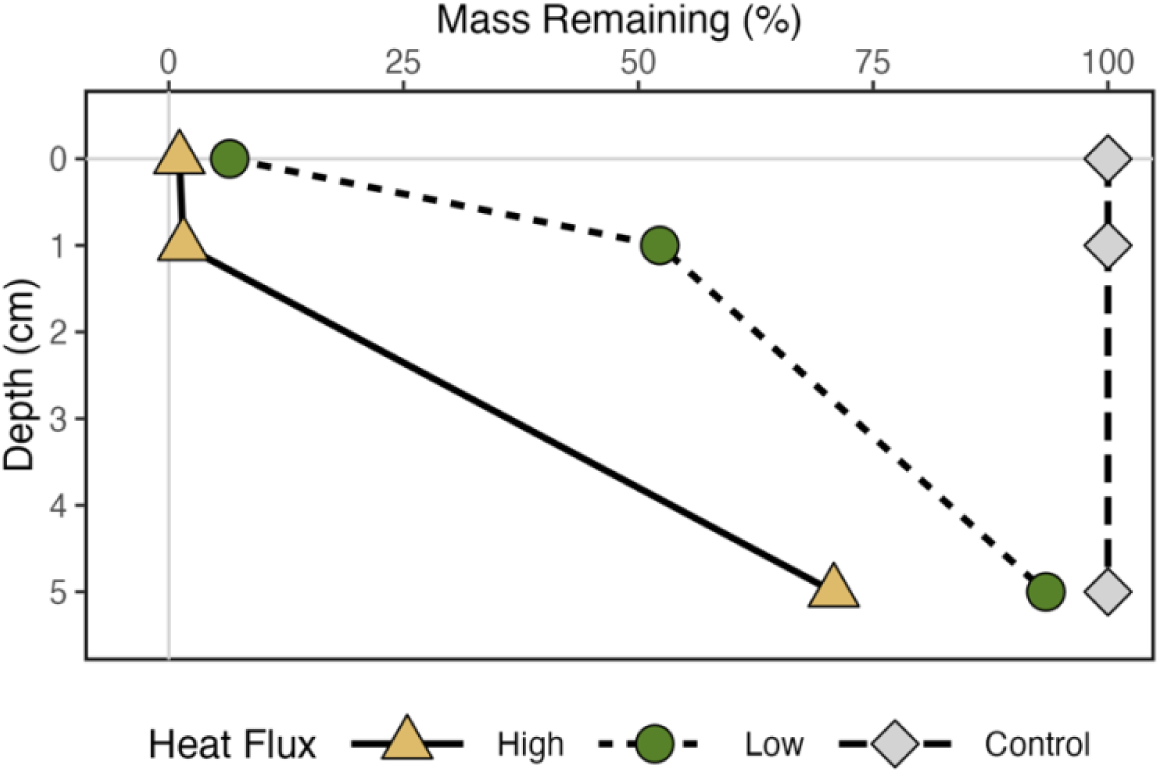
Percent of PyOM mass remaining after each treatment.

**Figure 9.**
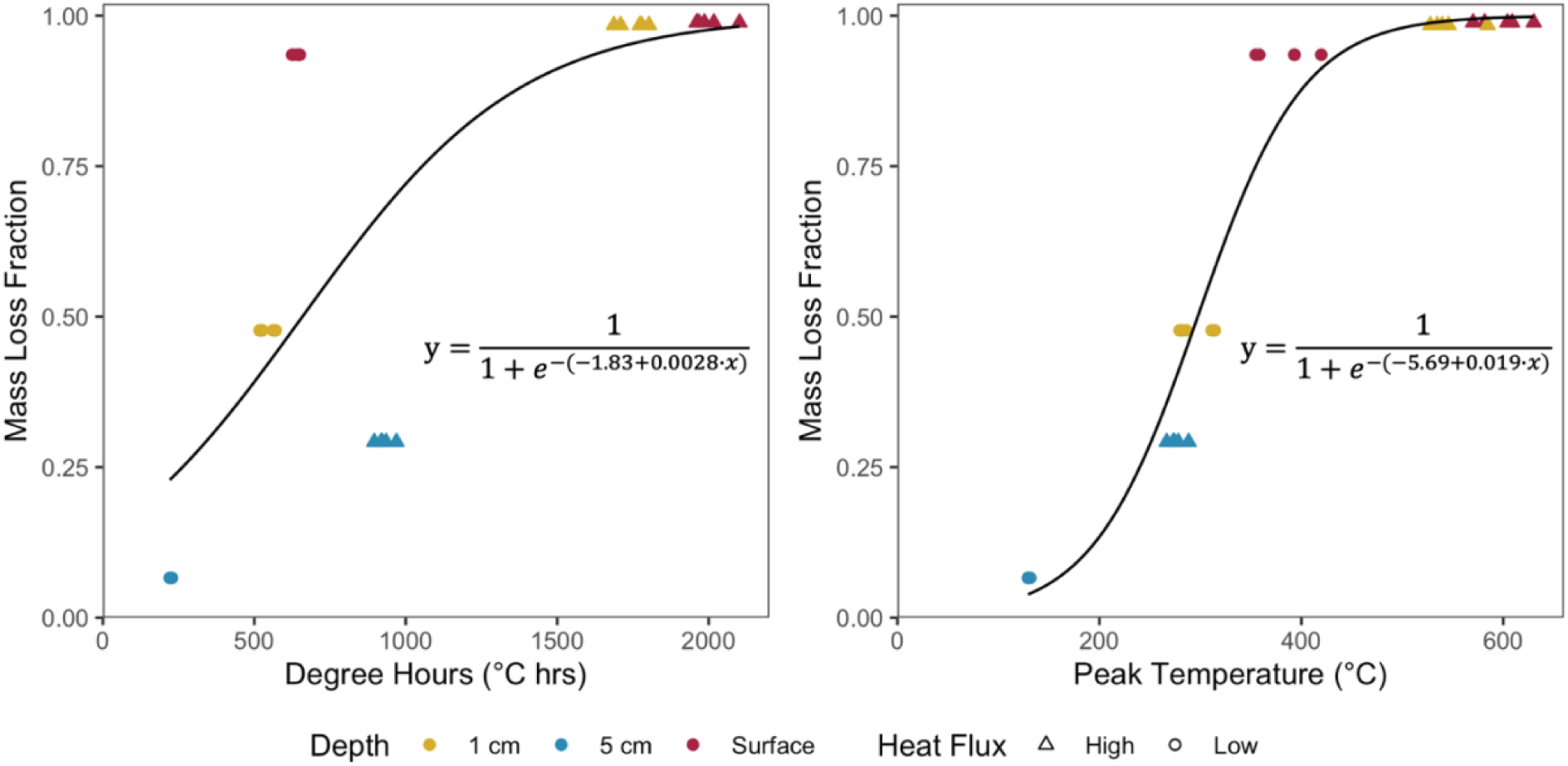
Relationship between degree hours (left; curve indicates logistic model fit; p < 0.05 for all coefficients; pseudo-R^2^=0.05; AIC = 31.316) or peak temperature (right; curve indicates logistic model fit; p < 0.05 for all coefficients; pseudo-R^2^=0.18; AIC = 18.311) and mass loss fraction

## Discussion

### Highly repeatable laboratory burns

The High- and Low-HF profiles created distinctive differences in degree hours, peak temperatures, and PyOM mass loss. The burn unit setup resulted in very consistent heat fluxes at a given depth, as indicated by the low variation among thermocouples located at samples within the same treatment. With little variation in the heat profiles dosed to the samples and different effects created by different treatments, this method should be considered promising and repeatable. Although the HF received by the burn unit is not directly monitored by HF sensors during the experimental burns, we could still ensure the HF-temperature conversion was stable throughout the burn. After the experimental burns were all completed, we conducted sand-only burns again under temperature control mode with only the HF sensor to test if there was any deviation in the HF that the samples received. The results from the last blank burns showed that the heat flux was typically within ±4 kW m^-2^ of the prescribed HF profile (Fig. S3 and S4).

### Interpretation of the system and potential limitations

Using quartz sand as the medium to contain the PyOM samples does not only ensure chemical and physical uniformity, but also simulates the soil texture in typical jack pine barrens. Because quartz has a high melting point (∼1700°C), is chemically inert even during extensive heating, and was ashed and washed before simulations, changes to the PyOM should be the only factor accounting for the difference in chemical properties for different treatments. This approach may be limited for finer soil textures and different mineralogy, where soil texture can change due to the fusion of clay minerals during heating (Badía and Martí 2003). Furthermore, such effects would complicate the heat transfer. In systems where using quartz sand is appropriate, we can ensure a relatively consistent heat transfer due to uniform particle size.

Some moisture would be naturally present in soil, even though the soil of jack pine barrens is sandy and well drained. However, moisture was not incorporated into the experiments for simplification purposes. In general, we would predict that a moist sand matrix would have higher heat capacity due to the presence of water, so it would take more energy for the temperature to increase than the dry sand matrix. Massman (2012) modeled sand heating and moisture transportation and indicated that moisture (14% of the volume) at and above 35 mm in a sand matrix evaporated at 100°C within the first hour of heating, and the temperature surged after the water was evaporated. We might thus predict that the near-surface treatments in our experiments would not be dramatically different in a moist system, but for samples at 5 cm, lower temperatures due to high moisture might decrease mass losses. An additional consideration is that water can also help create anoxic or low-oxygen (O_2_) conditions that slow down the oxidation of PyOM during heating. Although the O_2_ content in the burn unit was not directly measured, the general burn conditions for surface treatments were assumed to be aerobic. In theory, when surface heating is sufficient to initiate combustion >200°C, gas products tend to move upwards and exit the burn unit, and the replenishment of O_2_ at deeper soil profiles is much slower than its depletion (Bryant et al. 2005). Thus, we should likely assume that the samples in the buried treatments have experienced both aerobic and anaerobic heating.

### Even buried PyOM can still be combusted in subsequent fires

Consistent with our hypotheses, higher HF and shallower depths had more PyOM mass losses. Previous lab and field experiments by Doerr et al. (2018) and Bartoli et al. (2021) determined that low-intensity fires consumed 17-50% of PyOM in mass at the litter surface and soil surface, while high-intensity fires consumed 50-84%. In our study, the mass loss of PyOM at the surface was substantially higher – 93.51% under the Low-HF profile (equivalent to low-intensity fire) and 98.86% under the High-HF profile (equivalent to high-intensity fire). Our buried PyOM had substantial mass loss (>25%) for all buried treatments, except for 5 cm depth under Low-HF profile. This likely reflects the fact that our study was designed to represent the upper end of heat fluxes characteristic of realistic surface burns. Therefore, our results may overestimate total PyOM mass loss across a typical burn on a landscape scale but, rather, represent local conditions where logs are burning on the forest floor.

In wildland fires, fire may move quickly on a landscape due to various factors such as wind and topography, so the heating duration may be shorter than the HF profiles we applied here. For example, at spots where the fuel density is high, the fire may dwell longer. However, heating duration may not be a key determinant of PyOM mass loss after a certain amount of time, as we found that ‘degree hours’ was a poorer predictor of mass loss than peak temperature (Fig. 9).

Tinkham et al. (2016) indicated that PyOM is easily degraded by subsequent fires when on the surface or shallow-buried, while thermal degradation is less likely in mineral layers (>30 mm) due to soil insulation. The exposure depths in our study were designed to interrogate this concept. However, substantial thermal degradation still occurred at the depth of 5 cm, particularly under High HF. This can be caused by extensive low-temperature heating. Low-temperature heating (< 300°C) can also increase C degradability (Norwood et al. 2013; Santín et al. 2016a). Most carbonaceous material can start to be combusted at 150°C ∼ 250°C (Baldock and Smernik 2002; Badía and Martí 2003), which is also observed in our experiment. Due to its heterogeneous nature, it is not possible to determine an exact temperature required for PyOM to be thermally degraded. Based on our results, there is even mass loss for PyOM at 5 cm under low HF, which had a peak temperature of less than 150°C (Table 1). For all other treatments, the temperature stayed above 150°C for an extensive time (Fig. 8), which can create substantial thermal oxidation. Based on the logistic model fitting, we found that substantial mass loss of PyOM occurred at the peak temperature of 200 to 400°C, and stayed basically unchanged for the higher temperatures (due to near-complete combustion). Consequently, we could predict that PyOM even at 5 cm depth in soil profiles can be susceptible to consumption in subsequent fires, as long as the temperature reaches the limits required for thermal degradation, as a recent study has also found substantial decrease in soil mass and soil C after a high-severity wildfire (McCool et al. 2023).

## Conclusion

We presented detailed methods for simulating the effects of reburns on PyOM using HF parameterizations with log burns and applying the HF profiles to PyOM using a cone calorimeter. The methods we designed for HF simulation are highly replicable and provide distinctive temperature profiles and heat exposure for each treatment (with shallower depths and higher HFs resulting in higher heat exposure), which could be adopted for further soil heating experiments. We found that deeper burial depths can help protect PyOM from loss in the subsequent fires, and burial is a more effective protector under low-intensity fire. In future work, chemical and biological properties of the PyOM collected from the burn unit will be assessed using different lab test to further analyze the net effects of fire.

## Acknowledgement

We would like to thank Roger Bohringer from Wilson State Forest Nursery and Stuart Seaborne from Hancock Agricultural Research Station for the donation of jack pine trees. We also want to thank Troy Humphrey from Department of Soil Science for helping cut and grind the tree branches to produce PyOM and Kelsey Kruger for additional support and help.

## Data Availability Statement

Data presented in the paper will be open to access online through the US Department of Energy’s ESS-DIVE repository.

## Supplementary Information

**Table S1.** ANOVA statistics for peak temperature (°C)

| Sources | df | Sum of Squares | Mean Squares | F statistic | p-value |
| --- | --- | --- | --- | --- | --- |
| Heat Flux | 1 | 312785 | 312785 | 879.48 | $< 2 \times 10^{-16}$ |
| Depth | 2 | 451137 | 225568 | 634.25 | $< 2 \times 10^{-16}$ |
| Heat Flux × Depth | 2 | 14462 | 7231 | 20.33 | $6.83 \times 10^{-6}$ |
| Residuals | 24 | 8536 | 356 |  |  |

**Table S2.** ANOVA statistics for degree hours (°C × hrs)

| Sources | df | Sum of Squares | Mean Squares | F statistic | p-value |
| --- | --- | --- | --- | --- | --- |
| Heat Flux | 1 | 8976887 | 8976887 | 7449.7 | $< 2 \times 10^{-16}$ |
| Depth | 2 | 3040982 | 1520491 | 1261.8 | $< 2 \times 10^{-16}$ |
| Heat Flux × Depth | 2 | 600963 | 300481 | 249.4 | $< 2 \times 10^{-16}$ |
| Residuals | 24 | 28920 | 1205 |  |  |

**Figure S1.**
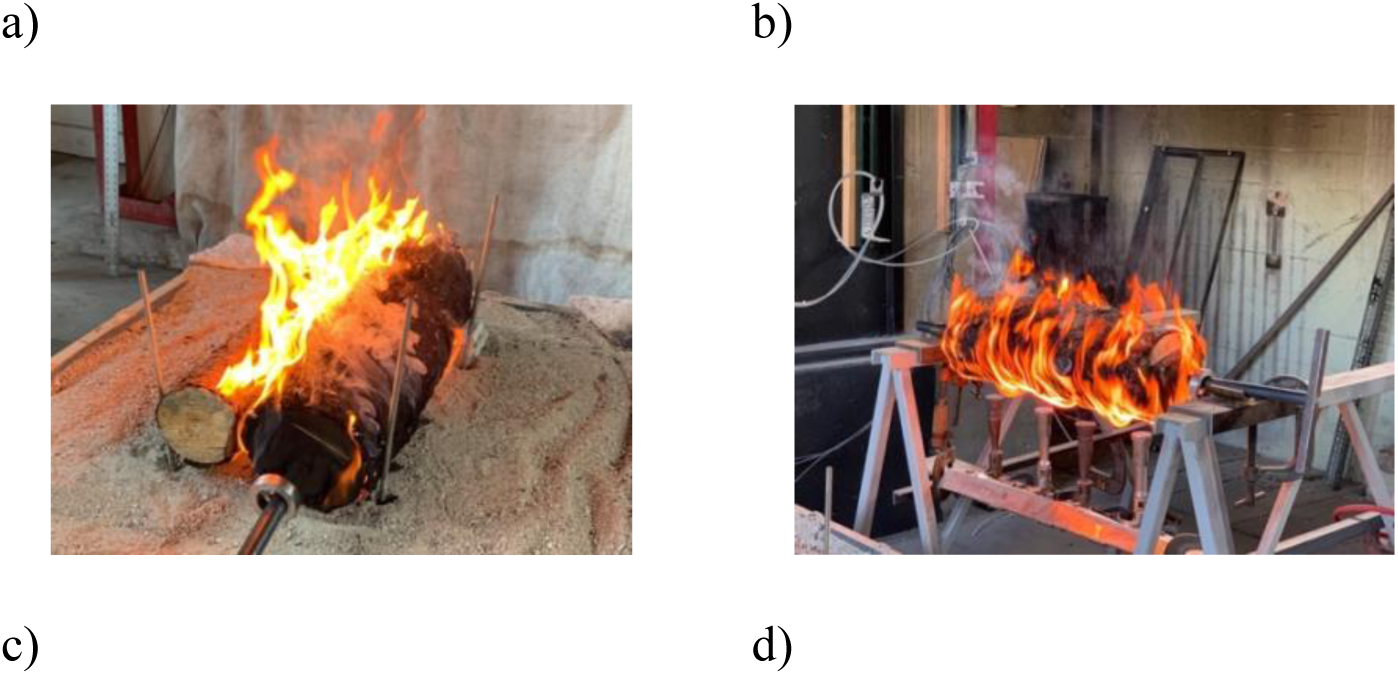
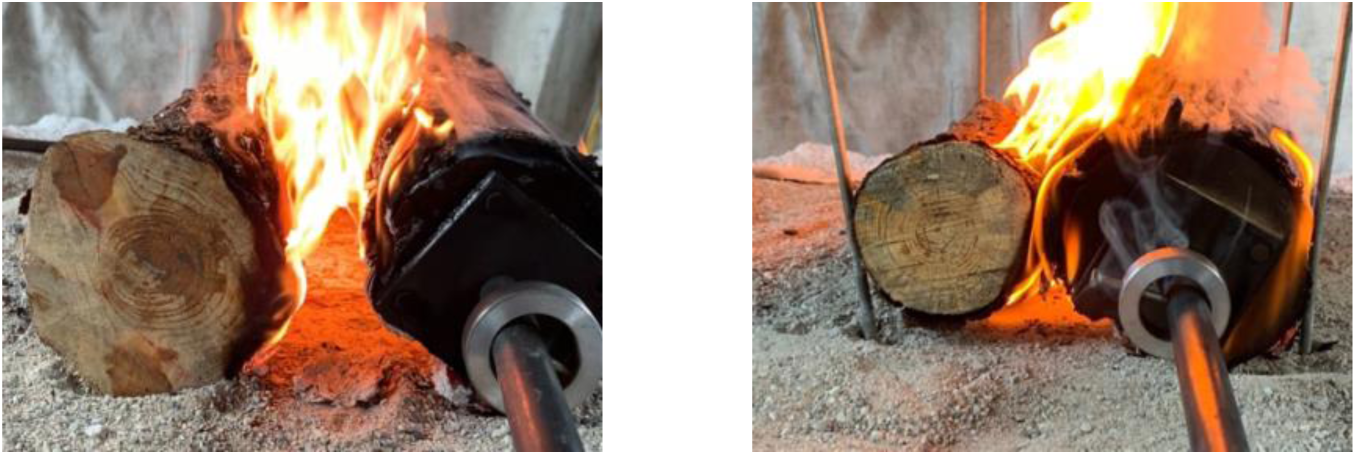
Log burns preparation and setup. a) Logs and sensors’ placement; b) Treated log on burners; c) the treated (right) and untreated (left) logs on the sand bed for capturing Low HF profile; d) the treated (right) and untreated (left) logs on the sand bed for capturing High HF profile

**Figure S2.**
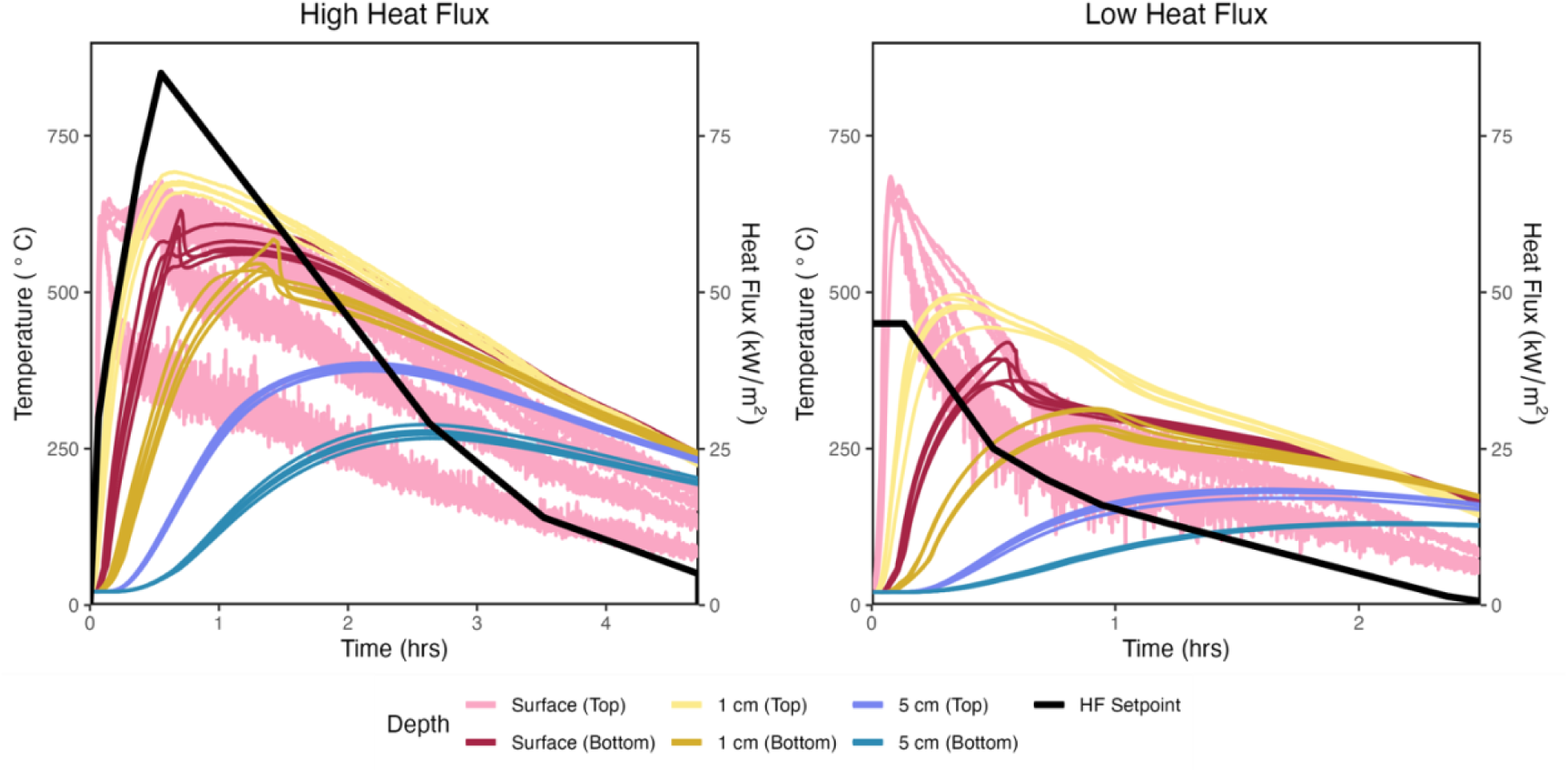
Temperature profiles for top and bottom thermocouples for five replicate samples in each HF treatment (colored lines) and the corresponding heat flux profiles (black lines) for High HF (left) and Low HF (right). Note different time scales on x-axis. Samples continued to cool to room temperature beyond data plotted here.

**Figure S3.**
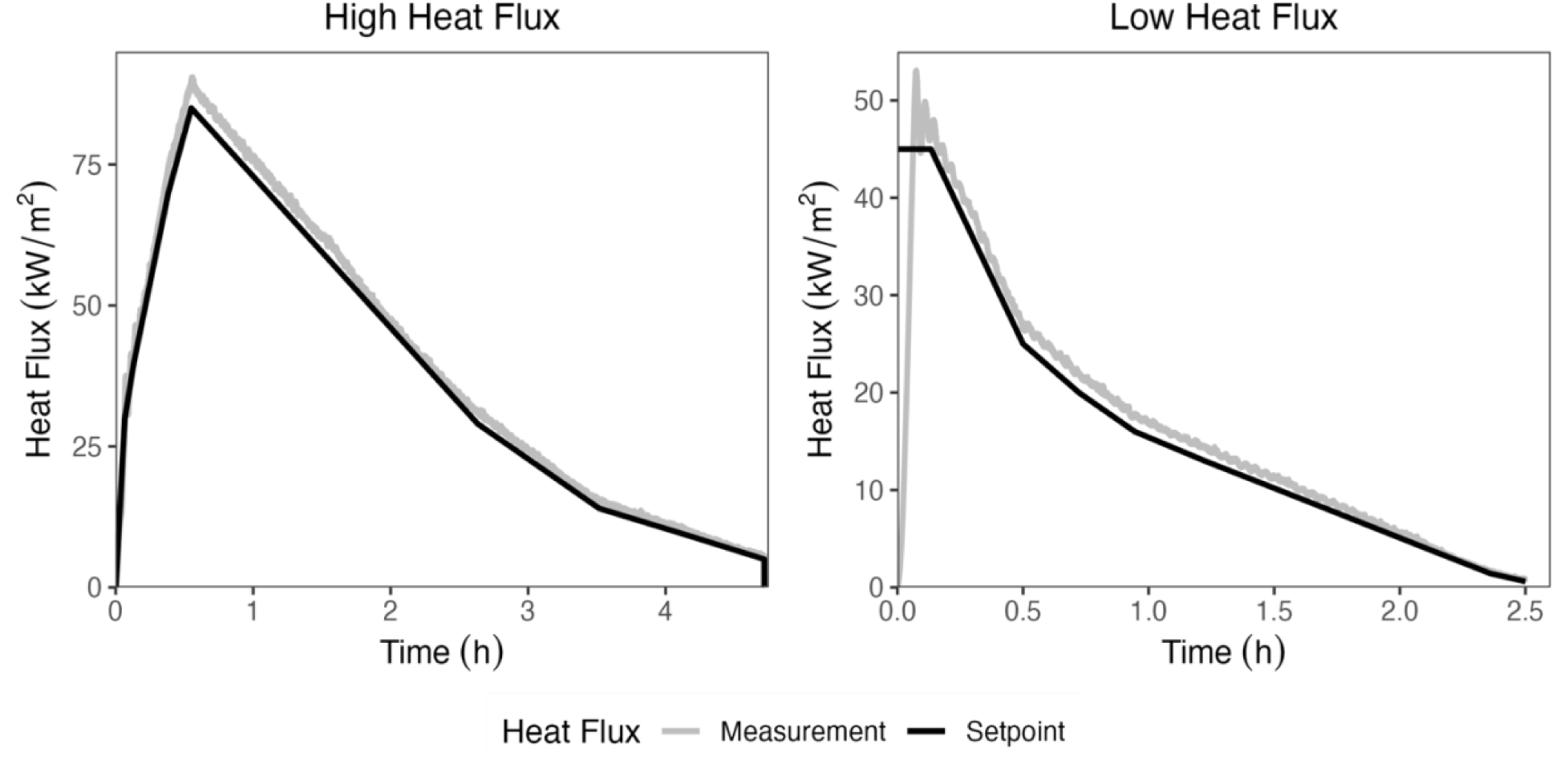
Comparison of the heat flux profiles (black line) with the heat flux detected by the sensor during a sand-only burn under temperature control mode (grey line) for High HF (left) and Low HF (right). Note different time scales on x-axis.

**Figure S4.**
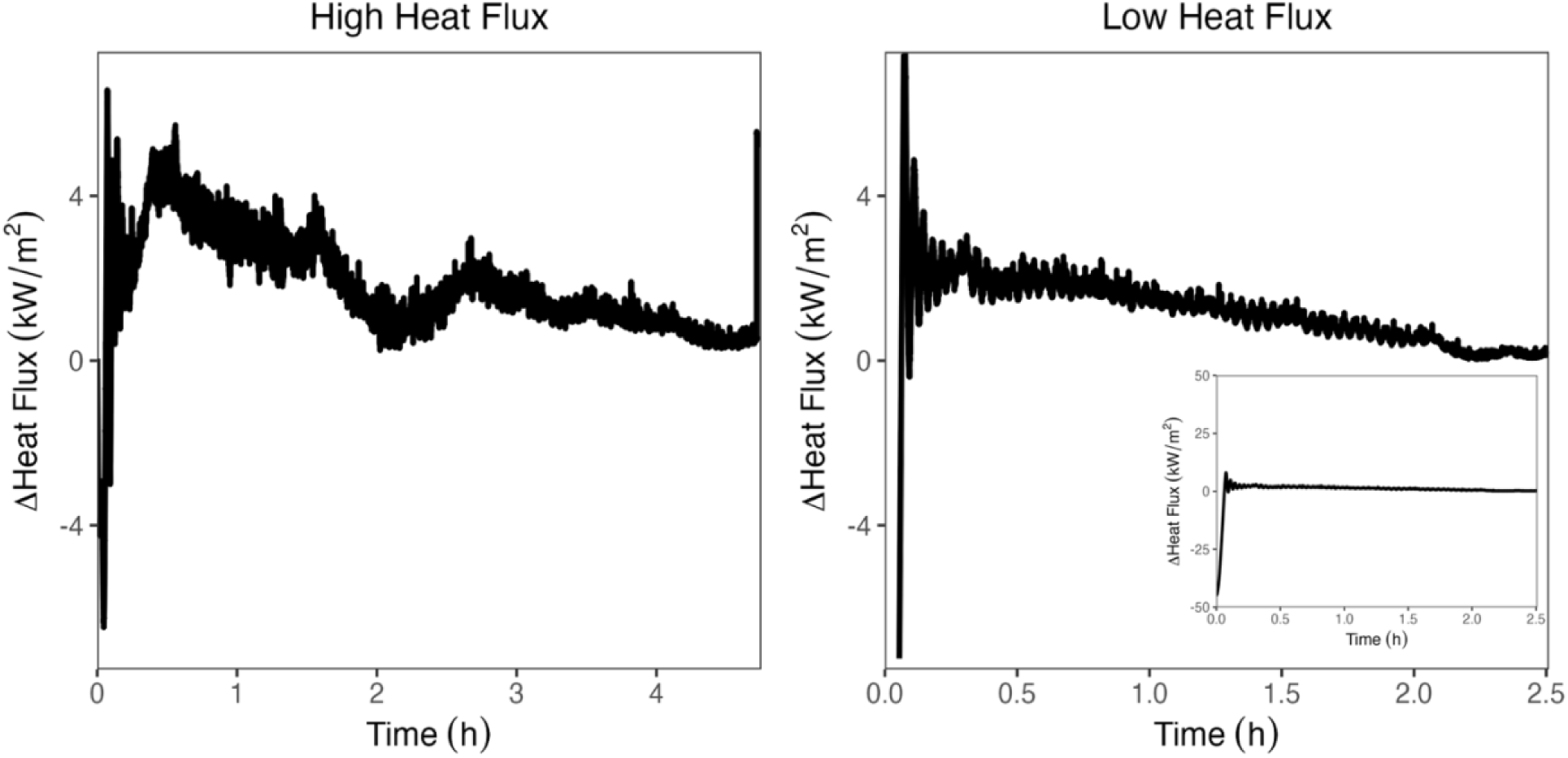
Difference between heat flux profiles with the heat flux detected by the sensor during the sand-only burn under temperature control mode for High HF (left) and Low HF (right). ΔHeat Flux (kW/m^2^) = Measured heat flux – Heat flux setpoint.

